## supplemental information for "Functional repertoire convergence of distantly related eukaryotic plankton lineages revealed by genome-resolved metagenomics"

### Supplementary material

#### Chapters:

#1: Genome-resolved metagenomics with anvi'o

#2: Decontamination of single cell genomes with anvi'o

#3: The METdb database for eukaryotic transcriptomes

#4: Categorizing the 939 TARA Oceans metagenomes

#5: Manual curation of the DNA-dependent RNA polymerase genes (SMAGs and METdb)

#6: World map projections

#### Genome-resolved metagenomics with anvi'o

##### # A set of single copy core genes to identify eukaryotic MAGs

As initially outlined in a blog post published at the beginning of this project to benefit others[1], we have defined a set of 83 single copy core genes from BUSCO[2] compatible with the gene calling workflow of anvi'o[3] to best estimate the completion of eukaryotic metagenome-assembled genomes (MAGs). Figure 1 describes the efficacy of this collection to estimate completion of MAGs from *Micromonas* and *Ostreococcus*. Note that those estimates are only initial, since this stage of the workflow uses a gene calling (Prodigal[4]) that is not optimal for eukaryotes. However, the results are sufficiently robust to effectively guide the manual binning and curation of eukaryotic MAGs without the need to first identify eukaryotic contigs in the assembly output. While the identification of eukaryotic contigs prior to binning as been benchmarked by the group of Jill banfield[5], false positives and false negatives associated with this critical step can be problematic and are entirely avoided in our workflow. We found that binning metagenomes containing multiple domains of life can be done smoothly within anvi'o, as long as proper single copy core gene collections are used to efficiently affiliate MAGs to Bacteria, Archaea and Eukarya. Note that this dedicated collection for eukaryotes is the main improvement within anvi'o compared to the workflow outlined for the characterization of ~1,000 bacterial and archaeal MAGs from small size fractions of TARA Oceans[6]. It is now an integral component of the anvi'o metagenomic flow used by a growing number of scientists interested in genome-resolved metagenomics.

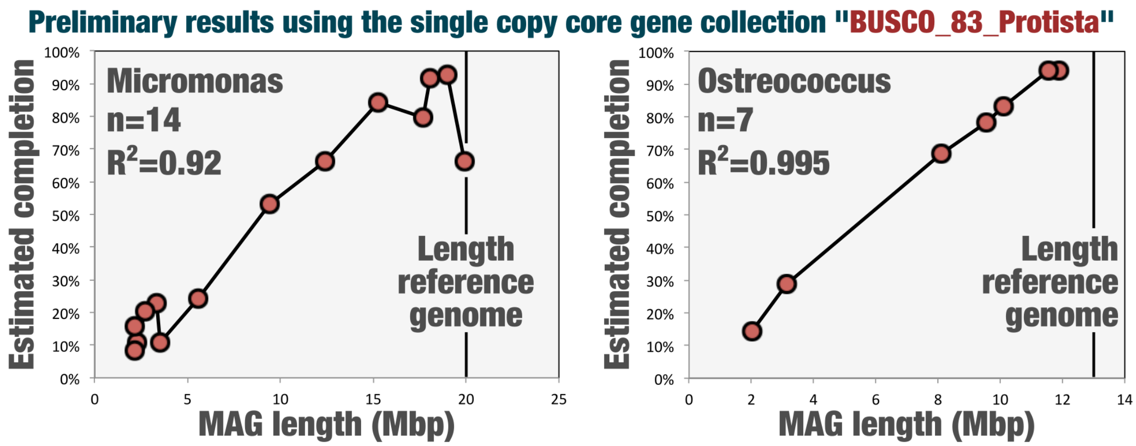


**Figure 1: Completion estimates for *Micromonas* and *Ostreococcus* MAGs using a set of 83 BUSCO single copy core genes, as a function of the length of the MAGs.**

#### # A summary of the workflow to bin and curate eukaryotic MAGs

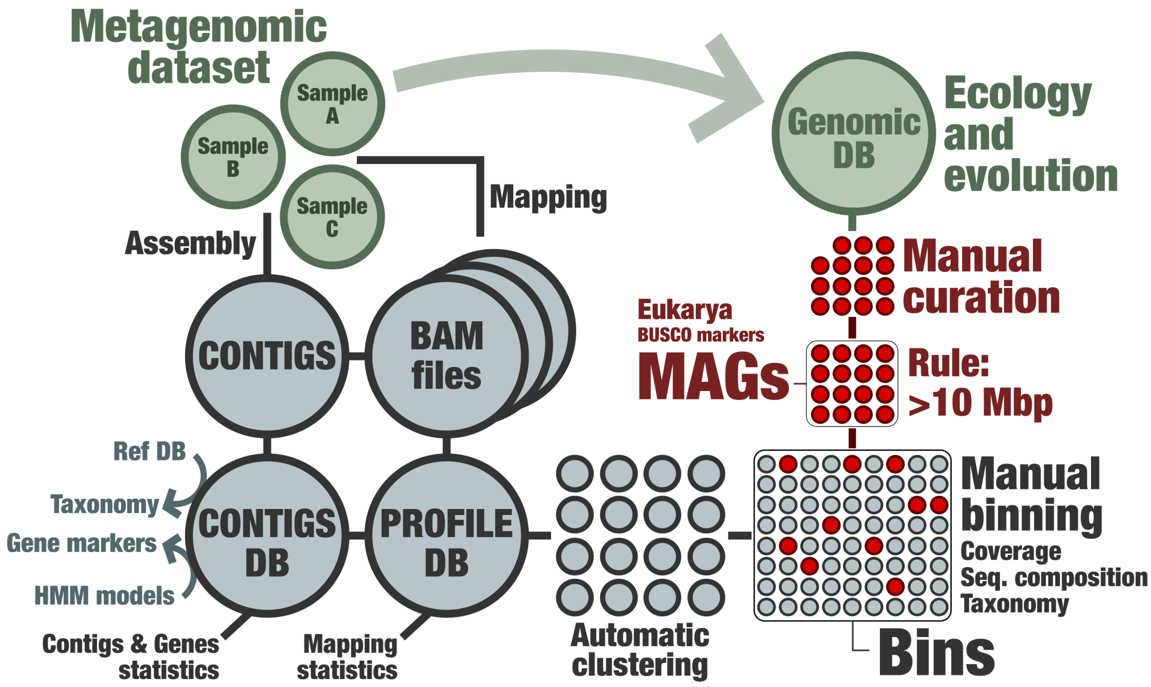


**Figure 2: The manual genome-resolved metagenomic framework of anvi'o dedicated to the eukaryotes. This workflow is to be applied to each assembly outcome.**

We followed the workflow outlined in the figure 2 for each of the 11 metagenomic co-assemblies outlined in the study (see Table S2). Briefly, we used the sequence composition of contigs and their differential coverage across metagenomes to perform a first automatic binning step with CONCOCT[7] by constraining the number of created clusters to a number substantially below the number of genomes in the assembly. This number ranged from 50 to 400 depending on the assembly volume. Note that CONCOCT is used because the interactive interface of anvi'o cannot work efficiently when loading >25k contigs. For each of the CONCOCT clusters, we then used the anvi'o interactive interface to manually identify and curate eukaryotic MAGs. This step took about 10 months of manual work.

### An holistic interactive interface now compatible with eukaryotes

Within the framework of our study, the anvi'o interactive interface took advantage of the sequence composition of contigs, their differential coverage across metagenomes, taxonomic signal using a reference database that includes METdb, and HMM models for single copy core gene collections (Bacteria, Archaea, Eukarya). When selecting a cluster of contigs corresponding to a MAG in the interface, anvi'o identified its domain affiliation in real time using random forest, and displayed its completion and redundancy values accordingly. This way, it was possible to focus on the eukaryotic MAGs within an assembly containing also many abundant bacterial and archaeal MAGs.  In the figure 3, we provide the example of one CONCOCT cluster from the Mediterranean Sea metagenomic co-assembly (95 metagenomes) containing eukaryotic MAGs for *Ostreococcus* and *Micromonas* (left panel). In this simple example, we selected those two clusters in the interface, saved the collection, and subsequently manually curated them as presented here for *Ostreococcus* (right panel). This MAG exhibited a completion of 100% and a redundancy of 3%. One metagenome (most outer blue layer) was particularly useful in this particular case since the *Micromonas* MAG was more detected compared to the Ostreococcus MAG, allowing an effective binning outcome. Given the complexity of marine metagenomes, differential coverage across dozens of metagenomes strongly benefited to the outcome of our genome-resolved metagenomic survey.

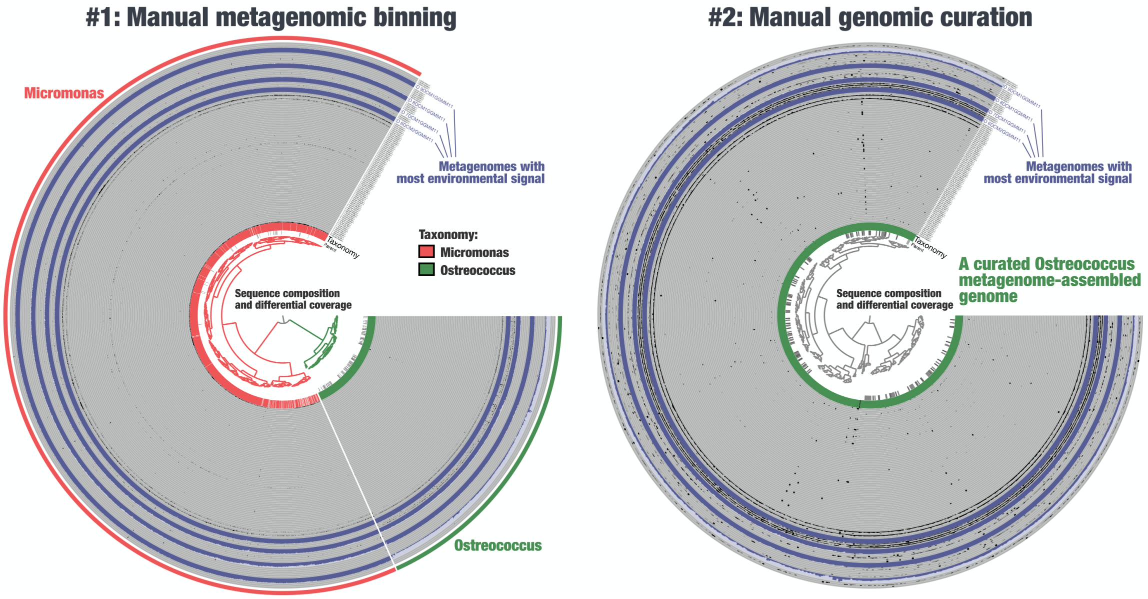


**Figure 3: The anvi'o interactive interface to manually bin and curate eukaryotic MAGs. The left panel displays the detection of contigs from a single CONCOCT cluster across 95 metagenomes, alongside taxonomic signal. Clustering was done using sequence composition and differential coverage. Right panel displays the curated *Ostreococcus* MAGs identified from the left panel.**

### Decontamination of single cell genomes with anvi'o

Eukaryotic single cell genomes (SAGs) can be heavily contaminated due to a combination of factors during cell sorting, DNA extraction and amplification, and multiplex sequencing. Here, we slightly modified the anvi'o metagenomic workflow to effectively decontaminate marine eukaryotic SAGs, one by one. Briefly, we used the anvi'o interactive interface to manually curate eukaryotic SAGs by taking into consideration the sequence composition of contigs, their differential coverage across 100 most relevant metagenomes (i.e., those with highest mapping recruitment scores within the scope of TARA Oceans), taxonomic signal using a reference database that includes METdb, and HMM models for single copy core gene collections (Bacteria, Archaea, Eukarya). Note that compared to the metagenomic co-assemblies, the number of contigs under consideration was orders of magnitude smaller. Since all contigs could be loaded in the interactive interface, there was no need to use the pre-clustering step with CONCOCT. However, CONCOCT could also be used here if some SAG assemblies include more than ~25k contigs.

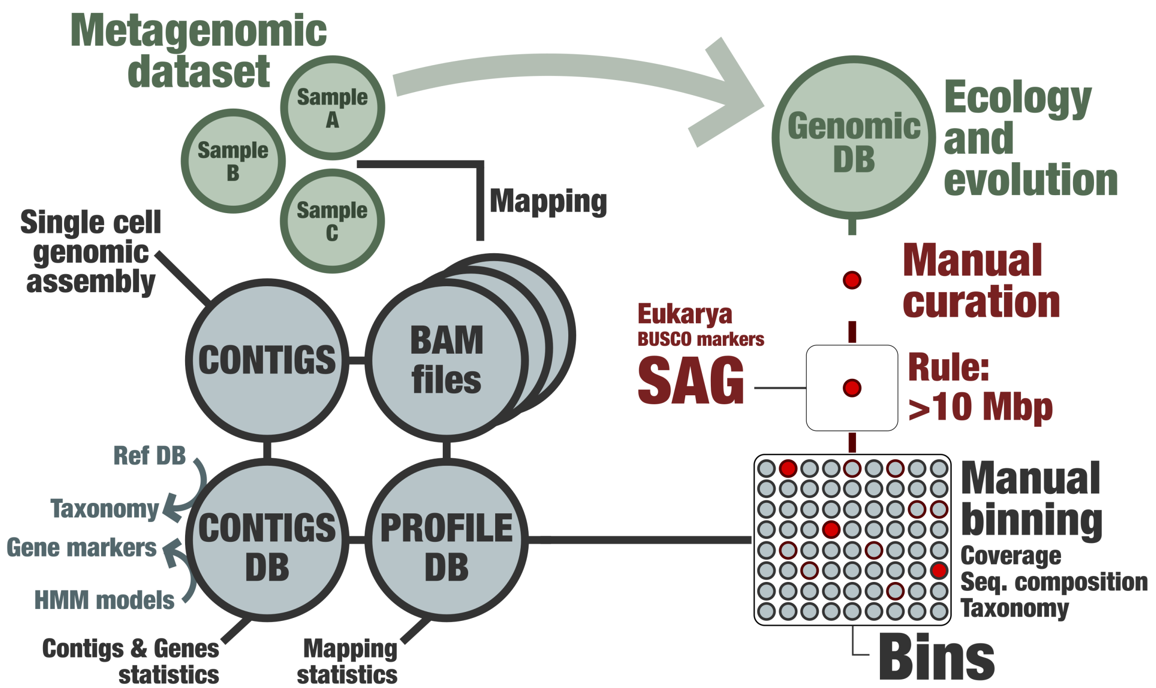


**Figure 5: The manual metagenomic framework of anvi'o dedicated to the decontamination of SAGs. This workflow is to be applied to each SAG assembly outcome.**

Figure 6 provides a striking example of heavily contaminated SAG we could effectively curate thanks to the clear differential coverage signal of contigs across 100 metagenomes. In this particular case, contamination seemed to have multiple origins, and a large number of contigs were removed. Overall, our manual curation of SAGs using a genome-resolved metagenomics workflow initially built for MAGs turned out to be highly valuable, leading in our study to the removal of more than one hundred thousand scaffolds for a total volume of 193.1 million nucleotides. This metagenomic-guided decontamination effort contributes to previous efforts characterizing eukaryotic SAGs from the same cell sorting material[8–12] and provides new guidelines for marine eukaryotic SAGs. This approach is now highly recommended for future efforts generating eukaryotic SAGs from the sunlit ocean. This is important, especially since SAGs could become a valuable asset in the near future to target lineages genome-resolved metagenomics failed to recover so far. It is especially the case of Dinoflagellates.

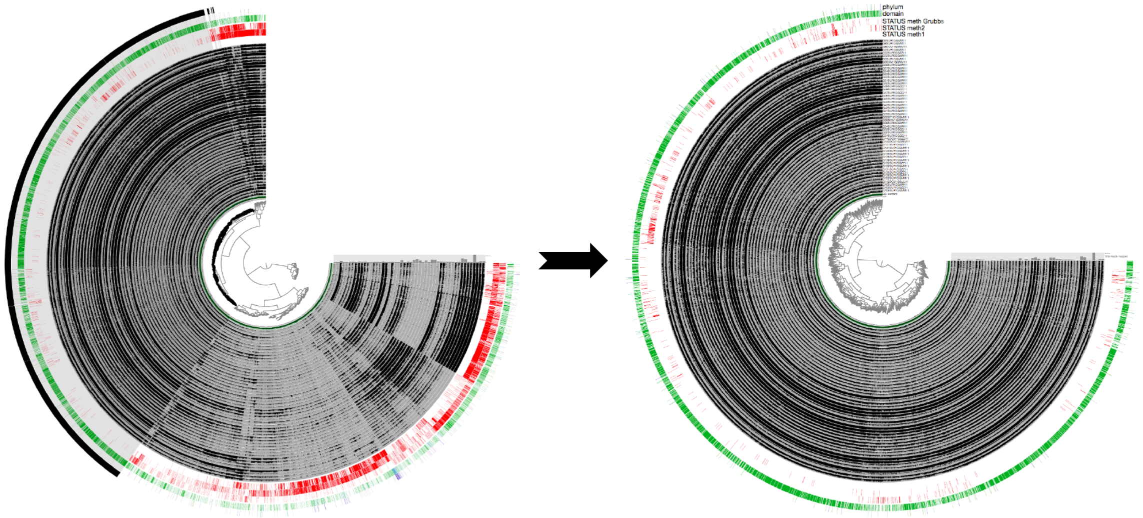

**Figure 6: Example of the decontamination of TOSAG00-8. Left panel describes all contigs reconstructed from this SAG. The selection of contigs (outer layer) corresponds to our final curated SAG, displayed in the right panel.**

### The METdb database for eukaryotic transcriptomes

METdb is a curated database of transcriptomes from marine eukaryotic isolates that cover the MMETSP collection[13] (new assemblies were performed, combining time points from the same culture in co-assemblies when available) as well as cultures from TARA Oceans. The associated manuscript is not yet published. However, the database is publically available and can be accessed at http://metdb.sb-roscoff.fr/metdb/.

### Categorizing the 939 TARA Oceans metagenomes

Our study surveyed a total of 939 TARA Oceans metagenomes (Table S1) that we organized into four cellular size categories (**size 1**: 0.2-5µm, **size 2**: 3-20µm, **size 3**: 20-200µm, **size 4**: 180-2000µm) as well as a wider cellular size fraction encompassing all categories considered in our study (**wider size**: 0.8-2000µm). The four cellular size categories were well represented across the five oceans and two seas. Overall, 119 stations contained at least 3 out of the 4 cellular size categories, which we defined as **Station subset 1** (757 metagenomes). Using this first subset,

SMAGs were assigned a "**cosmopolitan score**" corresponding to the percentage of stations in which they were detected. SMAGs were also assigned a "**cellular size range"** and "**oceanic signal"** using average coverage in each size categories (n=4) for the former and in each ocean and sea (n=7) for the later. Those results are summarized in the tables S3 an S4. Unfortunately, the wider cellular size fraction was missing in the Mediterranean Sea, Red Sea and Indian Ocean, limiting its use to 91 stations from the four remaining oceans, which we defined as **Station subset 2** (130 metagenomes). Critically, this second subset offers a glimpse into the relative proportion of planktonic lineages of different cellular sizes. While more limited in its geographic scope, the **Station subset 2** could provide important insights into the **"relative proportion**" of SMAGs in stations from the Atlantic, Pacific, Arctic and Southern Ocean.

### Manual curation of the DNA-dependent RNA polymerase genes for SMAGs and METdb

An eukaryotic dataset (Da Cunha)[14] was used to build HMM profiles for the two largest subunits of the DNA-dependent RNA polymerase (RNAP-a and RNAP-b). These two HMM profiles were incorporated within the anvi'o framework to identify RNAP-a and RNAP-b genes (Prodigal[4] annotation) in the SMAGs and METdb transcriptomes.

We independently performed the following workflow for RNAP-a sequences identified in the SMAGs (round A, n= 1,626) and METdb (round B, n= 2,823) as well as for RNAP-b sequences identified in the SMAGs (round C, n= 1,373) and METdb (round D, n= 3,941):

(1) **Stetting the stage with references:** Reference sequences for the relevant largest subunits of the DNA-dependent RNA polymerase (e.g., RNAP-a for round A) corresponding to eukaryotic (types I, II and III), bacterial and archaeal lineages from the Da Cunha dataset were added to the sequences identified by the HMM.

(2) **Phylogenetic tree Phase 1:** Sequences were aligned using the iterative FFT-NS-i refinement method of MAFFT[15] v7.464 with default parameters, and the sites with more than 50% of gaps were trimmed using Goalign v0.3.0-alpha5. Phylogenetic trees were reconstructed with IQ-TREE[16] v1.6.12. The model of evolution was estimated with the ModelFinder Plus option[17], and supports were computed from 1,000 replicates for the Shimodaira-Hasegawa (SH)-like approximation likelihood ratio (aLRT)[18] and ultrafast bootstrap approximation (UFBoot)[19]. Anvi'o v6.1 was used to visualize and root the phylogenetic trees.

(3) **Identifying sequences of type I, II and III:** We used the anvi'o interactive interface to root the tree between Bacteria and the rest, and identify sequences corresponding to eukaryotic DNA-dependent RNA polymerase of

type I, II and III. Sequences not clearly belonging to one of these three clusters were discarded. Note that during this process other types of eukaryotic RNA polymerase (e.g., nucleomorphs) were identified and put aside for investigations beyond the scope of this study.

(4) **Fusing fragmented sequences when needed:** For each SMAG or METdb transcriptome, sequences corresponding to the same RNA polymerase type (e.g., RNAP-a_type_I for round A) were aligned against each other and against a relevant eukaryotic reference sequence using blastp[20]. Non-overlapping sequences corresponding to the same subunit (based on **Phylogenetic tree Phase 1**) were considered fragments of the same gene and fused manually, overcoming fragmentation issues during gene calling and/or transcription. In addition, only the longest sequence was kept for overlapping isoforms and closely related duplicates (>95% identity and >30% coverage).

(5) **Phylogenetic tree Phase 2:** A phylogenetic tree was performed for each subunit (DNA-dependent RNA polymerase of type I, II and III) as done for the **Phylogenetic tree Phase 1** (for improved resolution, archaeal references were used as outgroup and bacterial sequences removed in this analysis). Distantly related duplicates (those occurred in <5% of SMAGs and <10% of METdb transcriptomes, possibly due to contamination) were carefully considered in the context of the three phylogenetic trees as well as taxonomy to identify and remove sequences with incoherent phylogenetic and/or taxonomic signal.

(6) **Final collection:** We removed sequences shorter than 200 amino-acids, providing a final collection of DNA-dependent RNA polymerase genes for the SMAGs (n=2,150) and METdb (n=2,032) with no duplicates.

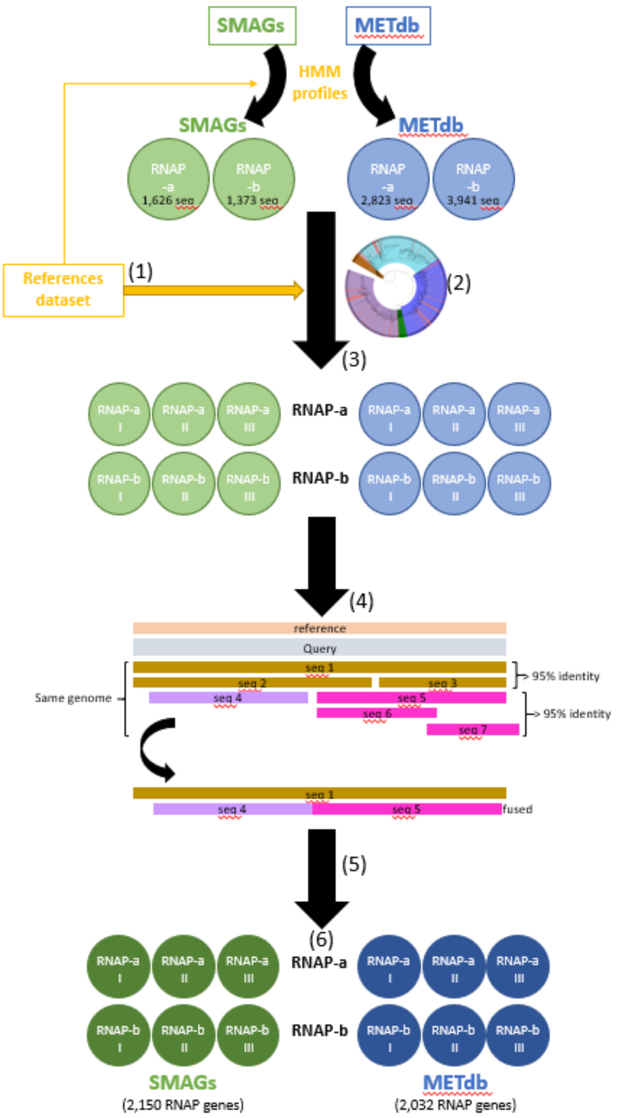


### World map projections

#### # World Ocean Atlas data

Seven physicochemical parameters were used to define environmental niches: sea surface temperature (SST), salinity (Sal), dissolved silica (Si), nitrate ($NO_3$), phosphate ($PO_4$), iron (Fe), and a seasonality index of nitrate (SI $NO_3$). With the exception of Fe and SI NO3, these parameters were extracted from the gridded World Ocean Atlas 2013 (WOA13)[21]. Climatological Fe fields were provided by the biogeochemical model PISCES-v2[22]. The seasonality index of nitrate was defined as the range of nitrate concentration in one grid cell divided by the maximum range encountered in WOA13 at the Tara sampling stations. All parameters were co-located with the corresponding stations and extracted at the month corresponding to the Tara sampling. To compensate for missing physicochemical samples in the Tara *in situ* data set, climatological data (WOA) were favored. The correlation between *in situ* samples and corresponding values extracted from WOA were high:

**# R-squared values for the surface samples:**
SST: 0.99, Sal: 0.86, Si: 0.89, NO3: 0.85, PO4: 0.90

**# R-squared values for the DCM samples:**
SST: 0.97, Sal: 0.47, Si: 0.97, NO3: 0.74, PO4: 0.85

In the absence of corresponding WOA data, a search was done within 2° around the sampling location and values found within this square were averaged.

Nutrients, such as $NO_3$ and $PO_4$, displayed a strong collinearity when averaged over the global ocean (correlation of 0.95 in WOA13), which could complicate disentangling their respective contribution to niche definition. However, observations and experimental data allow distinguishing between limiting nutrients at regional scale characterized by specific plankton communities[23]. The future projection of niches will yield spurious results when the present-day collinearity is not maintained[24,25]. To this day, there is no evidence for large scale changes in global nutrient stoichiometry[26].

#### # Earth System Models and bias correction

Outputs from six Earth system models were used to project environmental niches under greenhouse gas emission scenario RCP8.5[27]:

| Model | Reference |
|---|---|
| CESM1-BGC | Gent et al., 2011 |
| GFDL-ESM2G | Dunne et al., 2013 |
| GFDL-ESM2M | Dunne et al., 2013 |
| HadGEM2-ES | Collins et al., 2011 |
| IPSL-CM5A-LR | Dufresne et al., 2013 |
| IPSL-CM5A-MR | Dufresne et al., 2013 |
| MPI-ESM-LR | Giorgetta et al., 2013 |
| MPI-ESM-MR | Giorgetta et al., 2013 |
| NorESM1-ME | Bentsen et al., 2013 |

Environmental drivers were extracted for present day (2006-2015) and end of century (2090-2099) conditions for each model and the multi-model mean was computed. A bias correction method, the Cumulative Distribution Function transform, CDFt[28], was applied to adjust the distributions of SST, Sal, Si, $NO_3$ and $PO_4$ of the multi-model mean to the WOA database. CDFt is based on a quantile mapping (QM) approach to reduce the bias between modeled and observed data, while accounting for climate change. Therefore, CDFt does not rely on the stationary hypothesis and present and future distributions can be different. CDFt was applied on the global fields of the mean model simulations. By construction, CDFt preserved the ranks of the simulations to be corrected. Thus, the spatial structures of the model fields were preserved.

### Environmental niches models: training, validation and projections

From the initial dataset of 713 SMAGs, we selected those present in at least 4 stations for environmental niche training, discarding just 58 of them. Four machine learning methods were applied to compute environmental niches for each of the 655 remaining SMAGs:

(1) Gradient Boosting Machine (gbm)[29]
(2) Random Forest (rf)[30]
(3) Fully connected Neural Networks (nn)[31]
(4) Generalized Additive Models (gam)[32]

Hyper parameters of each technique (except gam) were optimized as followed:

(1) For gbm, the interaction depth (1, 3 and 5), learning rate (0.01, 0.001) and the minimum number of observations in a tree node (1 to 10)
(2) For rf, the number of trees (100 to 900 with step 200 and 1000 to 9000 with step 2000) and the number of parameters used for each tree (1 to 8)
(3) For nn, the number of layers of the network (1 to 10) and the decay (1.10-4 to 9.10-4 and 1.10-5 to 9.10-5)
(4) For gam the number of splines was set to 3.

R packages gbm (2.1.3), randomForest (4.6.14), mgcv (1.8.16) and nnet (7.3.12) were used for gbm, rf, nn and gam models.

To define the best combination of hyper parameters for each model, we perform 30 random cross-validations by training the model on 75% of the dataset randomly sampled and by calculating the Area Under the Curve51 (AUC) on the 25% remaining points of the dataset. The best combination of hyper parameters is the one for which the mean AUC over the 30 cross-validation is the highest. A model is considered valid if at least 3 out of the 4 techniques have a mean AUC superior to 0.65, which is the case for 374 out of the 655 SMAGs (57%). Final models are trained on the full dataset and only the techniques that have a mean AUC higher than 0.65 are considered to make the projections. The majority (286) of the 374 validated niches is validated by all four models and 88 by only 3 models. Relative influences of each parameter in defining environmental niches are calculated using the feature_importance function from the DALEX R package[33] for all four statistical methods. For model training and projections, physicochemical variables are scaled to have a mean of 0 and a variance of 1. For this scaling, the mean and standard deviation of each WOA13 variable (+ PISCES-v2 Fe) co-localized with Tara stations with a value available is used. This standardization procedure allows for better performance of models. Finally, as statistical models often disagree we use the ensemble model approach for global-scale projections of niches[34] i.e. the mean projections of the validated machine learning techniques.
